# Empowering Engineered Muscle Function by Extending Connexin 43 Duration with Reduced Graphene Oxides

**DOI:** 10.1101/2021.12.08.470989

**Authors:** Eunkyung Ko, Onur Aydin, Zhengwei Li, Lauren Gapinske, Kai-Yu Huang, Taher Saif, Rashid Bashir, Hyunjoon Kong

**Author notes:** Corresponding author Rashid Bashir, Ph.D., Dean, College of Engineering, Grainger Distinguished Chair in Engineering, Professor of Bioengineering, University of Illinois at Urbana-Champaign, 306 Engineering Hall, MC-266, 1308 W. Green Street, Urbana, IL 61801, (Email), Hyunjoon Kong, Ph.D., Robert W. Schafer Professor, Department of Chemical and Biomolecular Engineering, University of Illinois at Urbana-Champaign, Urbana, IL 61801-3602, (E-mail).

## Abstract

Engineered skeletal muscle act as therapeutics invaluable to treat injured or diseased muscle and a “living” material essential to assemble biological machinery. For normal development, skeletal myoblasts should express connexin 43, one of the gap junction proteins that promote myoblast fusion and myogenesis, during the early differentiation stage. However, myoblasts cultured in vitro often down-regulate connexin 43 before differentiation, limiting myogenesis and muscle contraction. This study demonstrates that tethering myoblasts with reduced graphene oxide (rGO) slows connexin 43 regression during early differentiation and increases myogenic mRNA synthesis. The whole RNA sequencing also confirms that the rGO on cells increases regulator genes for myogenesis, including troponin, while decreasing negative regulator genes. The resulting myotubes generated a three-fold larger contraction force than the rGO-free myotubes. Accordingly, a valveless biohybrid pump assembled with the rGO-tethered muscle increased the fluid velocity and flow rate considerably. The results of this study would provide an important foundation for developing physiologically relevant muscle and powering up biomachines that will be used for various bioscience studies and unexplored applications.

## INTRODUCTION

The development of a reliable *in vitro* skeletal muscle model system has the potential to benefit both fundamental and applied bioscience studies. For instance, skeletal muscle recreated with patients’ stem or precursor cells can simulate development, regeneration, and pathogenesis and screen newly developed therapeutics. The engineered muscle may also be transplanted to treat acute muscle injuries and chronic skeletal degenerative diseases.^[1]^ Recent studies have used engineered muscle to power biological machines capable of locomotion and performing desired functions.^[2, 3]^ Success relies on the ability to reproduce muscle with physiologically relevant contraction force. However, the engineered skeletal muscle generates the force usually in micronewton scale, which is smaller than in vivo muscle.

Various approaches have been made to enhance force generation by exercising the engineered muscle or adding growth factors or small molecules during maturation.^[4]^ Recently, the early loss of gap junction protein during myogenic differentiation has been brought to attention as a limiting factor. Gap junctions connect neighboring cells and allow intercellular transports of ions, electrical signals, and small metabolic molecules.^[5]^ Among the gap junction proteins, connexin 43 is abundant in cardiomyocytes and skeletal myoblasts.^[6, 7]^ The connexin 43 promotes intercellular association during myogenesis and propagates action potential signals through the cells, leading to synchronized muscle contraction.^[8, 9]^ It has also been reported that connexin 43 may contribute to muscle tissue regeneration.^[10]^ Conversely, suppression of connexin 43 in mice impact skeletal muscle development, as evidenced by reduced muscle weight.

Cardiomyocytes retain connexin 43 from the early to late differentiation stage. Skeletal myoblasts also present connexin 43 during the early differentiation in vivo.^[8, 10]^ However, they undergo downregulation after cells fuse to form myotubes. Then, motor neurons innervating myotubes take over control of muscle contraction.^[7, 11]^ In contrary, skeletal myoblasts cultured in vitro undertake early connexin 43 downregulation even before myogenic differentiation initiates, leading to limited myofiber formation and contraction.

To resolve this challenge, this study hypothesizes that reduced graphene oxide (rGO) flakes tethered to myoblasts would extend the duration of connexin 43 and subsequently recreate muscle with increased contraction frequency and force. In particular, rGO anchored on cells would promote the adsorption of extracellular matrix (ECM) proteins, which would support the assembly of connexin 43 into gap junctions between cells. In this study, we tethered rGO to C2C12 skeletal myoblasts, a model muscle precursor cell (Figure 1A). The tethered cells were collected to evaluate myogenic differentiation both in 2D and 3D. We first examined the extent to which rGO sustains cellular connexin 43 expressions and enhances myogenic differentiation by conducting immunofluorescence staining and quantitative real-time polymerase chain reaction (qRT-PCR) at different stages of differentiation. We also performed the whole RNA-sequencing to observe transcriptional activities. Lastly, we tested if the extended connexin 43 expression powers up 3D muscle contraction using a biohybrid Pump-Bot. This biological robot was designed to recapitulates the blood circulation mechanism in an embryonic vertebrate heart.^[3]^

**Figure 1.**
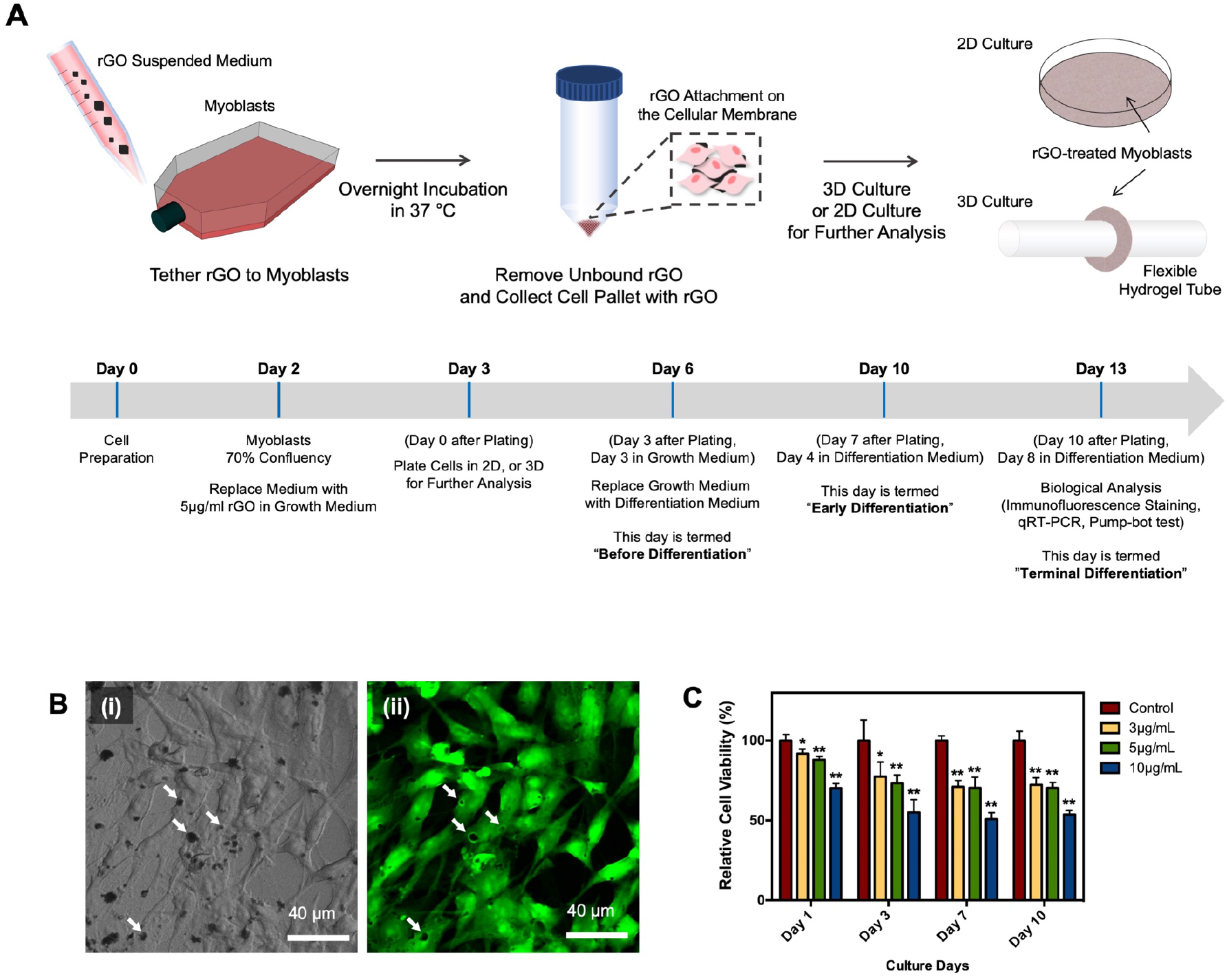
Tethering rGO flakes to skeletal myoblasts. (A) Procedure to prepare and analyze engineered muscle tethered with rGO. Myoblasts are first incubated in the rGO flakes-suspending medium overnight at 37 °C. The next day, cells tethered with rGO flakes are collected and used to prepare a monolayer cell sheet or a 3D muscle ring. The growth medium is replaced with a myogenic differentiation medium on day 3. After 7 days, the quality of engineered muscle was evaluated by examining genomic patterns, phenotypes, and contraction. **(B**) (i) Optical image of the myoblast tethered with rGO flakes. (ii) Fluorescently quenched rGO flakes on cells. In this image, the cell membrane was labeled with fluorescein. The white arrows indicate representative quenched areas. (C) Viability of myoblasts tethered with rGO using different concentrations of rGO-suspending media. Each bar value and error bar represent the average and standard deviation of four different samples per condition (*p < 0.01, **p < 0.05).

## RESULTS AND DISCUSSION

### rGO Flakes Adhere to Cell Surface with Minimal Internalization

First, we incubated myoblasts with rGO flakes with an average length of 5µm and width of 4µm, as shown with the transmission electron microscopy image (Figure S1A). Within 12hrs, rGO adhered to myoblasts stably. The Raman spectrum of the rGO on cells showed two distinct peaks, known as the *G* and *D* bands (Figure S1B). Centrifugation removed unbound rGO flakes while keeping rGO flakes tethered to the cell surface, as shown with brightfield images (Figure 1B(i)). We reconfirmed the rGO bound to the cell surface by photo-bleaching fluorescein associated with rGO (Figure 1B(ii)). According to the quantification made with photo-bleached images, about 1% of the cell surface was covered with rGO flakes. No flakes were found within cells, indicating that cells did not take up the micro-sized rGO flakes.

We also evaluated the cytotoxicity of rGO to myoblasts by incubating cells in the media, suspending controlled concentrations of rGO flakes, and counting the number of dead cells stained for Trypan blue (Figure 1C). More than 70% of cells remained viable during incubation with 3 and 5µg/ml rGO flake suspensions for 24hrs. According to the International Organization for Standardization, materials that keep cell viability at a level higher than 70% are considered biocompatible (10993-5).^[12]^ In contrast, the fraction of viable myoblasts incubated with 10µg/ml rGO was dropped to 70% within 24hrs. Therefore, the following studies kept the concentration of the rGO flakes in the incubation media at 5µg/ml. At this concentration, cells minimally internalized the micro-sized rGO flakes.

### rGO flakes Tethered to Myoblasts Extend Duration of Connexin 43 Expressions

Cellular connexin 43 expressions were analyzed by performing immunofluorescence imaging on day 3 (before differentiation), day 7 (early differentiation), and day 13 (terminal differentiation) (Figure 2A). We compared connexin 43 expressions of the rGO-tethered myoblasts with untreated ones. In both groups, connexin 43 was visible before differentiation. However, the untreated myoblasts lost connexin 43 expressions after they started differentiation. In contrast, rGO-tethered myoblasts continued to express connexin 43 during the early differentiation stage. The expression was finally downregulated at the terminal differentiation stage.

**Figure 2.**
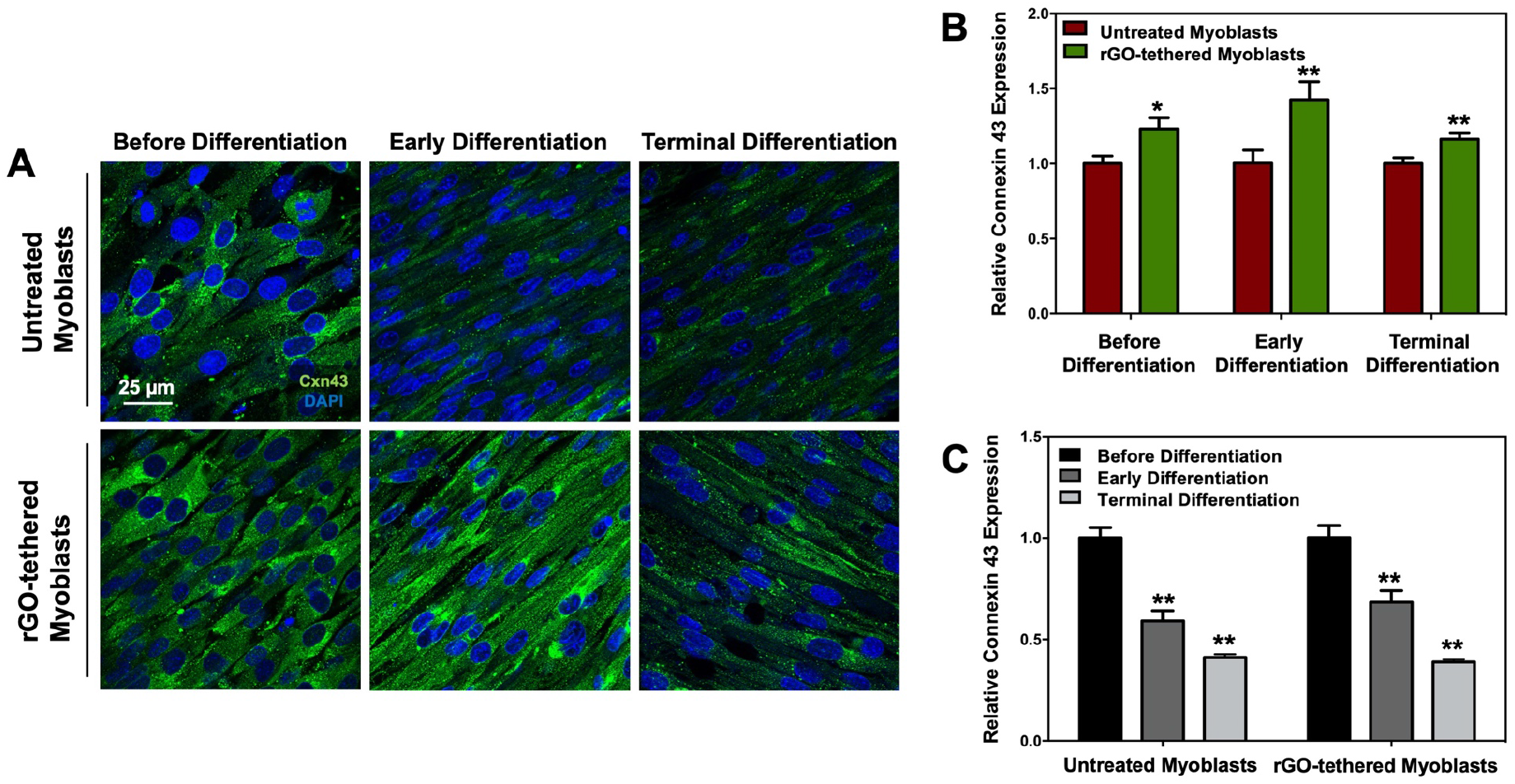
Time-dependent change of the connexin 43 expression in skeletal myoblasts. (A) Immunofluorescence images of connexin 43 (Cxn 43, green) and nucleus (DAPI, blue) in untreated and rGO-tethered cells. (B) Relative connexin 43 expression level of rGO-tethered myoblasts normalized to untreated myoblasts before differentiation (day 3), at early differentiation (day 7), and at terminal differentiation (day 13). (C) Change in relative connexin 43 mRNA expression level in the untreated and rGO-tethered myoblasts over time. The values were normalized to those measured before differentiation (day 3) for each group. In (B) and (C), each bar value and error bar represent the average value and standard deviation of three different samples per condition (*p < 0.01, **p < 0.05).

We additionally performed qRT-PCR to quantify the relative connexin 43 mRNA expression level. The connexin 43 mRNA expression level of rGO-tethered myoblasts was 1.2-fold higher on day 3, 1.4-fold higher on day 7, and 1.2-fold higher on day 13 than untreated group (Figure 2B). We also confirmed that connexin 43 mRNA expression was downregulated more rapidly in untreated myoblasts than rGO-tethered myoblasts (Figure 2C).

### rGO flakes Bound to Myoblasts Enhance Myotube Formation

We tested if the rGO flakes tethered to myoblasts enhance myogenic differentiation by performing immunofluorescence imaging and qRT-PCR. Cells were evaluated at 3 different time points; before differentiation (day 3), early differentiation (day 7), and terminal differentiation (day 13). The immunofluorescence images of myosin heavy chain showed that untreated cells form MF20-positive myotubes during the early differentiation stage (Figure 4A, first row). In contrast, rGO-tethered myoblasts developed MF20-positive myotubes before differentiation started (Figure 3A, second row). The myoblasts were also stained with F-actin and sarcomeric α-actinin (Figure 3B). Before differentiation, both untreated and rGO-tethered cells expressed F-actin only. As the differentiation proceeds, the multinucleated myotubes showed the sarcomeric α-actinins. Interestingly, the rGO-tethered cells formed larger myotubes with striated α-actinins than untreated cells.

**Figure 3.**
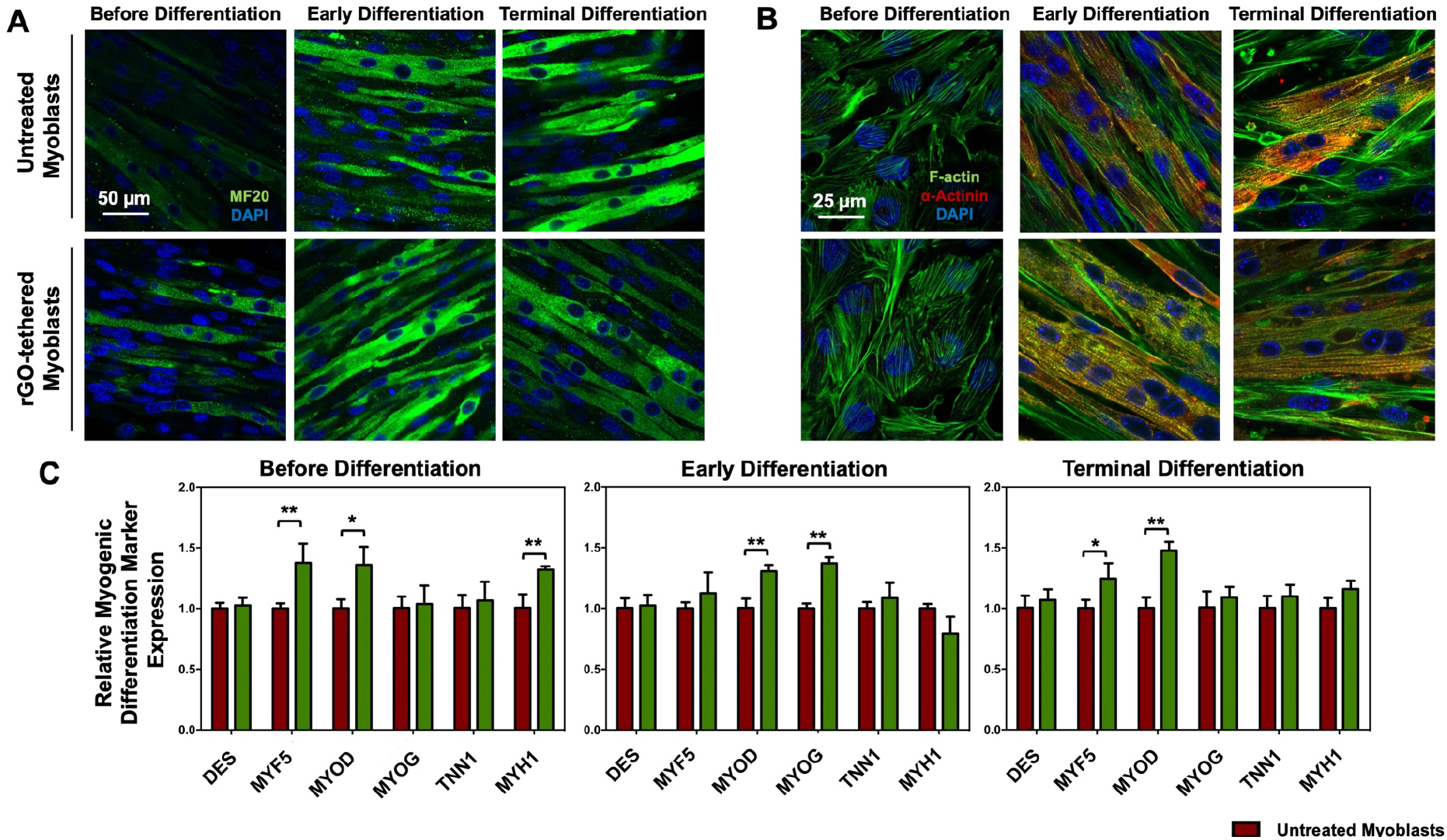
Effects of rGO on myogenic differentiation. (A) Immunofluorescently stained myotubes formed with untreated and rGO-tethered myoblasts. Myotubes were stained for myosin heavy chain (MF20, green) and nuclei (DAPI, blue). The images were captured before differentiation (day 3), at early differentiation (day 7), and at terminal differentiation (day 13). (B) Immunofluorescence images of myotubes stained for sarcomeric alpha-actinin (α-actinin, green) and nucleus (DAPI, blue). Both untreated and rGO-tethered cells were used to engineer myotubes. (C) Quantitative, real time-PCR analysis was performed to show relative myogenic marker gene expression levels in untreated and rGO-tethered myoblasts before differentiation (day 3), at early differentiation (day 7), and at terminal differentiation (day 13). The myogenic gene expression levels of rGO-tethered cells were normalized to that of untreated cells. Each bar value and error bar represent the average value and standard deviation of three different samples per condition (*p < 0.01, **p < 0.05).

**Figure 4.**
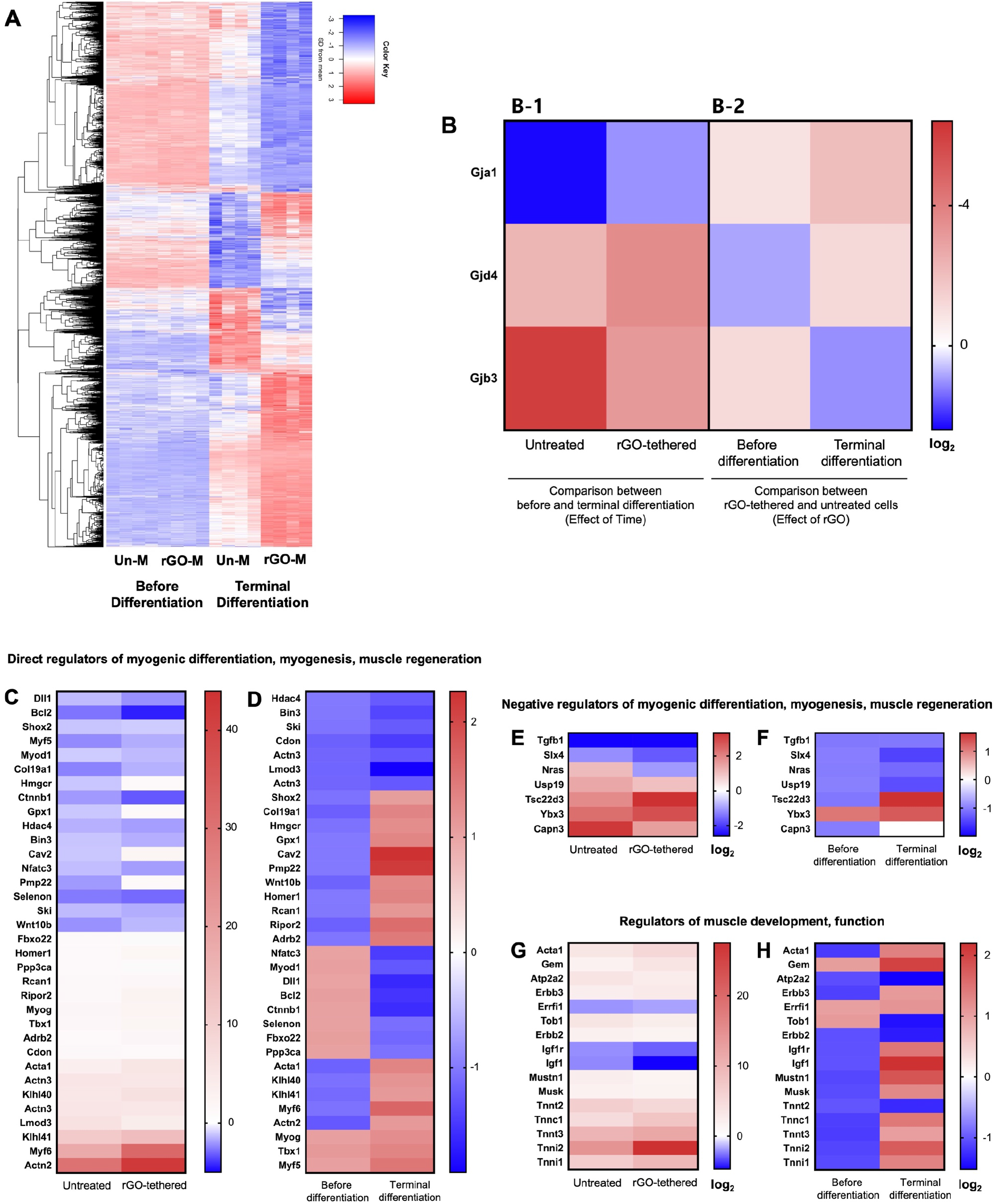
rGO-induced gene expression profile analysis by RNA sequencing. (A) Heat map showing the differentially expressed genes (total 7,474 genes, p-value<0.05, n=4). The color key indicates the gene expression level; red indicates increased expression, and blue indicates decreased expression level. (B) Gap junction-related gene expression change in untreated and rGO-tethered myoblasts (FDR p-value<0.05). (C-F) Quantification of gene expression changes that regulate myogenesis, myogenic differentiation, and muscle regeneration. The genes were selected from the heat map. (C) displays the positive myogenic regulatory gene expression change between day 3 (before differentiation) and day 13 (terminal differentiation) for untreated and rGO-tethered cells. (D) displays the positive myogenic regulatory gene expression difference between rGO-tethered and untreated cells before differentiation and at the termination stage. (E) displays the negative myogenic regulatory gene expression change between day 3 (before differentiation) and day 13 (terminal differentiation) for untreated and rGO-tethered cells. (F) displays the negative myogenic regulatory gene expression difference between rGO-tethered and untreated cells before differentiation and at the termination stage. (G) displays the muscle development and function-related gene expression change between day 3 (before differentiation) and day 13 (terminal differentiation) for untreated and rGO-tethered cells. (H) displays the muscle development and function-related gene expression difference between rGO-tethered and untreated cells before differentiation and at the termination stage. For each graph, the order of the genes is based on the fold increase of the first column.

These results illustrate that rGO flakes on myoblasts help to promote myogenic differentiation at the early stage. Given that rGO flakes serve to sustain cellular connexin 43 expressions at the early differentiation stage, the increased myogenic differentiation with rGO tethering is attributed to the extended duration of connexin 43. It is likely cells tethered by rGO flakes communicate better with adjacent cells through gap junctions, catalyzing the synthesis of myosin heavy chains.^[13]^

The myogenic differentiation was also evaluated by examining myogenic gene expression using qRT-PCR (Figure 4C). We compared mRNA expression levels of desmine (DES), myostatin (MSTN), myogenic factor 5 (MYF5), myogenic factor 5 (MYF5), myoblast determination protein (MYOD), myogenin (MYOG), troponin I (TNNI), and myosin heavy chain 1 (MHC1). These myogenic genes have been reported to regulate cell differentiation, myogenesis, and force generation.^[14, 15]^ Before initiation of differentiation (i.e., day 3), cells tethered with rGO flakes exhibited higher mRNA expressions for early myogenic proteins, including MYF5 (1.38 fold), MYOD (1.36 fold), and MYH1 (1.31 fold), than untreated cells. MYF5 and MYOD promote myogenic differentiation in an embryo, while MYH1 converts chemical energy to mechanical energy for muscle contraction. On day 7, rGO flakes on the cells increased MYOD (1.30 fold) and MYOG (1.37) gene expression levels. MYOG is another protein that promotes myogenic differentiation. The MYH1 gene expression was decreased slightly with rGO flakes, but no statistical significance of difference was found between conditions. On day 13, rGO flakes-tethered myoblasts showed a 1.24-fold increase in the MYF5 gene and a 1.47-fold increase in MYOD gene expressions, compared with untreated cells.

### rGO Flakes on the Myoblasts Regulate Cellular Genomics

We further examined the extent to which rGO regulates the whole genomic profile of myoblasts. Cells tethered with and without rGO were collected on day 3 (before differentiation) and day 13 (terminal differentiation stage) for analysis (Table 1). Before differentiation, untreated and rGO-tethered myoblasts expressed 160 different genes. At the terminal differentiation stage, the number of differentially expressed genes between two cell groups increased to 9,452.

**Table 1.** The number of differentially expressed genes analyzed with FDR p-value < 0.05. The effects of the rGO tethering and differentiation stage were separately analyzed to identify the number of upregulated and downregulated genes. Five contrasts were made to evaluate the effects of rGO, the effect of time, and lastly, the interaction (i.e., the time effect in the rGO group vs. the time effect in the control group).

|  | **Effect of Time**<br>(Before Differentiation vs Terminal Differentiation) |  | **Effect of rGO**<br>(Untreated Myoblasts vs rGO-tethered Myoblasts) |  | **Interaction** |
| --- | --- | --- | --- | --- | --- |
|  | Untreated myoblasts | rGO-tethered myoblasts | Before Differentiation | Terminal Differentiation |  |
| **Down Regulated** | 5440 | 5887 | 132 | 4769 | 3734 |
| **Up Regulated** | 5393 | 5887 | 28 | 4683 | 3740 |
| **Not Significant** | 3197 | 2256 | 13870 | 4578 | N/A |

Based on the interaction result, we compared 7,474 differentially expressed genes through differentiation with a p < 0.05 value and plotted those genes on a heatmap (Figure 4A). The color key shows if the gene was relatively upregulated (red) or downregulated (blue). First, we tracked gap junction-related genes in the heatmap (Figure 4B). Three genes include Gja1 (gap junction protein alpha 1) encoding connexin 43, Gjd4 (gap junction protein delta 4) encoding connexin 39, and Gjb3 (gap junction protein beta 3) encoding connexin 31. Both untreated and rGO-tethered myoblasts showed decrease in Gja1 expression after differentiation. The downregulation, however, was less significant with rGO-tethered myoblasts. The expression of the Gjd4 gene was increased in cells tethered with rGO, implying the increased expression of connexin 39, which regulates embryonic muscle development and fusion.^[16]^ The Gjb3 gene encoding connexin 31, not known to be expressed in skeletal muscle, was downregulated in rGO-tethered cells.^[17]^ Separately, we compared Gja1 gene expression of untreated cells with rGO-tethered cells (Figure 4B). The rGO-tethered cells showed higher Gja1 gene expression levels than untreated cells at both pre-differentiation and terminal differentiation stages. The fold increase compared with untreated cells was more significant at the terminal differentiation stage.

We analyzed the genes regulating myogenic differentiation, myogenesis, and muscle regeneration (Figure 4C, D). Untreated myoblasts exhibited up-regulation of 17 genes during differentiation. Interestingly, rGO tethered on the cells served to upregulate 11 genes, including Adrb2, Homer1, Rcan1, Ripor2, Tbx1, MyoG, Acta 1, Klhl40, Klhl41, Myf6, Actn2 (Figure 4C). In addition, 9 genes, including Ski, Selenon, Nfatc3, Bin3, Hdac4, Ctnnb1, Myod1, Bcl2, Dll1, were down-regulated in the rGO-tethered myoblasts.

We additionally analyzed the effect of rGO on the myogenic gene expression patterns at pre-differentiation (day 3) and terminal differentiation stages (day 13) (Figure 4D). Compared to the untreated cells, rGO-tethered myoblasts upregulated early myogenic markers, including Nfatc3, Myod1, Dll1, Bcl2, Ctnnb1, Selenon, Fbxo22, and Ppp3ca before differentiation.^[18]^ These early markers were down-regulated at the terminal differentiation stage. The rGO-tethered myoblasts also down-regulated terminal differentiation markers including Klhl40, Khlh 41, Myf6, Actn2, Acta1 at day 3, and upregulated them at day 13.^[19]^ We further mapped the expression profiles of negative regulators for myogenic differentiation, myogenesis, and muscle regeneration. We first tracked the gene expression change throughout myogenic differentiation (Figure 4E). Four out of seven negative regulators such as Slx4, Nras, Usp19, and Capn3 were down-regulated in rGO-tethered cells more significantly than untreated cells throughout differentiation (Figure 4E). We also examined the extent that rGO influenced negative gene expression before differentiation and at terminal differentiation. Five negative regulators, Tgfb1, Slx4, Nras, Usp19, and Capn3, were down-regulated at both time points (Figure 4F).

In addition, we analyzed genes that involve muscle development and function (Figure 4G, H). The rGO upregulated expressions of 4 genes encoding troponin subunits, troponin C, troponin T3, troponin I. Genes encoding actin and GTP-binding protein were also upregulated during differentiation (Figure 4G). Furthermore, genes encoding troponin T2, which plays a role in cardiac muscle contraction, were down-regulated in the rGO-tethered cells (Figure 4H). rGO also stimulated cellular integrin and extracellular matrix protein (ECM)-related gene expressions. In particular, 20 out of 39 genes showed greater increase over time with rGO tethered to the mhyoblasts (Supporting Information Figure 2). Additionally, 10 out of 39 genes and 21 out of 39 genes showed increase in presence of rGO at before differentiation and terminal differentiation stage, respectively (Supporting Information Figure 2).

Taken together, we suggest that these mRNA-Seq analyses disclose intracellular dynamic response to rGO on the cell surface during myogenic differentiation and myogenesis. During myogenic differentiation, multiple signaling pathways change in response to fusion and development.^[20]^ The rGO flakes did not alter gene expression patterns dramatically at the early differentiation stage but made significant changes at the terminal differentiation stage. This result may be attributed to the increase in the retention of ECM proteins on the cell surface. We have observed that tethering rGO to the myoblasts increased the number of extracellular matrix related gene expression levels greater in terminal differentiation stage (Supporting Information Figure 2). Based on our result, it may be concluded that the presence of rGO enhances the expression of these proteins and thus, facilitates the interaction between the cells and ECMs.

## CONCLUSION

In conclusion, this study demonstrates that extending the duration of connexin 43 on skeletal myoblasts significantly enhances the quality of muscle engineered in vitro. Tethering rGO flakes onto myoblasts circumvented early down-regulation of connexin 43 during myogenic differentiation. As a result, rGO enhanced myogenic differentiation, confirmed with the immunostaining, RT-PCR, and whole-genome profile analysis. The rGO-tethered myotubes also generated a three-fold larger contraction in a biohybrid pump than rGO-free myotubes, increasing both fluid velocity and flow rate. Combined with the genomic analysis, we suggest that the enhanced muscle contraction is related to the increased troponin synthesis in the myotubes. Overall, this study provides a foundation for improving the physiological activities of engineered muscle and further the performance of the biological machine. This study will significantly impact methods to recreate tissue and organoid platforms for fundamental and applied bioscience studies on development, disease, and new therapeutics.

## MATERIALS AND METHODS

### Characterization and analysis of rGO

The rGO flakes were purchased from ACS Materials (ACS Material LLC, CA). The rGO flakes were characterized by the Raman spectroscopy (Raman-11, Nanophoton) that uses argon laser operating at 514 nm as an excitation source. The morphology of rGO flakes was analyzed with images captured using transmission electron microscopy (TEM, JEOL 2100 TEM). The length and width of an individual rGO flake were measured using ImageJ.

### C2C12 skeletal myoblast culture

Cells plated on a 2D substrate were cultured in a growth medium consisting of Dulbecco’s modified Eagle medium (DMEM) (Corning Cellgro) supplemented with 10% (vol/vol) fetal bovine serum (FBS, Lonza), 1% (vol/vol) L-glutamine (Cellgro Mediatech), and 1% (vol/vol) penicillin-streptomycin (PS, Gibco). For myogenic differentiation, the growth medium was replaced with a differentiation medium (DMEM supplemented with 10% horse serum, 1% PS, 1% L-glutamine and IGF-1) after day 3 in the growth medium. The cells were incubated in the differentiation medium for additional 11 days. Both growth and differentiation medium were supplemented with the fibrinolytic inhibitor aminocaproic acid (ACA, 1mg/ml).

### Tethering rGO to cells

The stock of 5 mg/ml rGO suspended in ethanol was prepared. The stock suspension was pipetted well and sterilized under UV for 30 min. Then, the stock suspension was vortexed at maximum speed for 10 minutes and sonicated in a bath sonicator for an additional 30 minutes. The stock solution was diluted in the growth medium at a concentration of 5 µg/ml. The myoblasts, which reached 80% confluency, were incubated with rGO overnight at 37 °C. The next day, cells tethered with rGO were collected by trypsinizing the cells from the cell culture flask. The cells were centrifuged to remove excess rGO for further use.

### Analysis of rGO tethered on cells

rGO tethered on the cell was visualized by using a quenching.^[23]^ Cells tethered with rGO were plated on a glass-bottom dish with a growth medium. The coating solution was prepared by adding 1 mg of fluorescein sodium salt powder in 10mL of 0.5 wt % polyvinylpyrrolidone (PVP, Sigma-Aldrich, Mw = 360,000)/ethanol solution. 100 µL of the coating solution was dispensed on the glass area and spun for 5 seconds at 500 rpm and 45 seconds at 4,000 rpm. The quenched area was visualized by exposing cells to the 488 nm Argon laser and imaging them with confocal microscopy (Zeiss LSM 700).

### Cytotoxicity test

Trypan blue exclusion assay was used to determine the number of viable cells after the rGO tethering. Myoblasts were incubated with 0 µg/ml, 3 µg/ml, 5 µg/ml, and 10 µg/ml of rGO flakes in the growth medium. The cells were collected on days 1, 3, 7, and 10. Then, the viable cells were counted after treating with trypan blue solution. Nine sections on the hemacytometer were counted, and a total of 4 samples were analyzed per condition each day to verify the result.

### Immunocytochemical Analysis

Differentiated cells were analyzed with immunostaining for connexin 43 and myotubes. Cells were fixed in 4% (w/v) paraformaldehyde (Sigma) for 30 min, permeablized with 0.1% (v/v) Triton X-100 (Sigma) for 5 min and incubated in a 2% goat serum for 45 min at room temperature. After completing each step, the samples were washed with PBS 2 times. After blocking, cells were incubated with primary antibodies at 4 °C overnight. To stain for connexin 43, cells were incubated with anti-connexin43 (GJA1 antibody, Abcam, Cambridge, U.K.). To stain for myogenic markers, another set of cells were incubated in a solution of MF-20 anti-MHC (1:400) (iT FX, Developmental Studies Hybridoma Bank, The University of Iowa Department of Biology) or a solution of anti-sarcomeric α-actinin antibody (Abcam, Cambridge, U.K.). On the next day, cells were washed with PBS twice. Then, anti-connexin43 was labeled with Alexa Fluor-488 donkey anti-rabbit IgG (1:500; Invitrogen). Cells incubated with MF-20 were treated with Alexa Fluor-488 goat anti-mouse IgG (1:500; Invitrogen), and those incubated with anti-sarcomeric α-actinin antibody were treated with Alexa Fluor-568 donkey anti-rabbit IgG (1:500) (Invitrogen) and fluorescein (FITC)-conjugated Phalloidin. All samples were incubated with secondary antibodies for 1 hour. Finally, the nuclei were labeled with 4’,6-Diamidino-2-Phenylindole (DAPI, Sigma). The fluorescence images were obtained with a multiphoton confocal microscope (LSM 710, Carl Zeiss).

### Quantitative real-time polymerase chain reaction analysis

Quantitative real-time polymerase chain reaction (qRT-PCR) analysis was conducted at 3 different time points; before differentiation (day 3 after seeding), early differentiation (day 7 after seeding), and terminal differentiation (day 10 after seeding). Note that the growth medium was replaced with the differentiation medium on day 3. Total RNA from the samples was extracted from cells using the RNeasy Mini Kit (Qiagen) following the manufacturer’s protocol. After extracting the mRNA, cDNA synthesis was performed with qScript cDNA Super Mix (Quanta Biosciences) from 100 ng of RNA, and the reaction was performed according to the manufacturer’s protocol. Then, SsoFast EvaGreen Supermix (Bio-Rad) was added to the cDNA and primers, and the mixtures were analyzed using CFX Connect Real-Time System (Bio-Rad). For the analysis, the cycle threshold (Ct) values were compared relative to the GAPDH and control samples.

### RNA sequencing analysis

The RNA samples were prepared with TruSeq Stranded mRNA-seq Sample Prep kits(Illumina). Differential gene expression analysis was performed using the limma-trend method on the logCPM values. Total 5 comparisons were made; (1) Before differentiation vs. Terminal differentiation of the untreated cells, (2) before differentiation vs. terminal differentiation of the rGO-tethered cells, (3) rGO-tethered cells vs. untreated cells before differentiation stage, (4) rGO-tethered cells vs. untreated cells at terminal differentiation stage, and (5) time effect for the rGO-tethered cells vs. time effect for the untreated cells. Multiple testing correction was done separately for each comparison using the False Discovery Rate (FDR) method. The differential expression was considered significant at FDR p-value < 0.05.

### Statistical Analysis

The statistical analyses were performed using unpaired Student’s t-test with Graph Pad Prism 6.0 (Graph Pad Software Inc., San Diego, CA, USA). Statistical differences were considered significant at a p-value smaller than 0.05.

## ACKNOWLEDGEMENT

This work was supported by the National Science Foundation (CBET-1932192 & STC-EBICS Grant CBET-0939511), the National Institute of Biomedical Imaging and Bioengineering of the National Institutes of Health (T32EB019944 to E. Ko), and the Korean Institute of Science and Technology-Europe. The content is solely the authors’ responsibility and does not necessarily represent the official views of the National Institutes of Health. We want to thank Dr. Alvaro G. Hernández, Dr. Chris L. Wright, and Dr. Christopher J. Fields from the Roy J. Carver Biotechnology Center for the help in producing RNA Sequencing libraries.

## SUPPORTING INFORMATION

**Supporting Information Figure S1.**
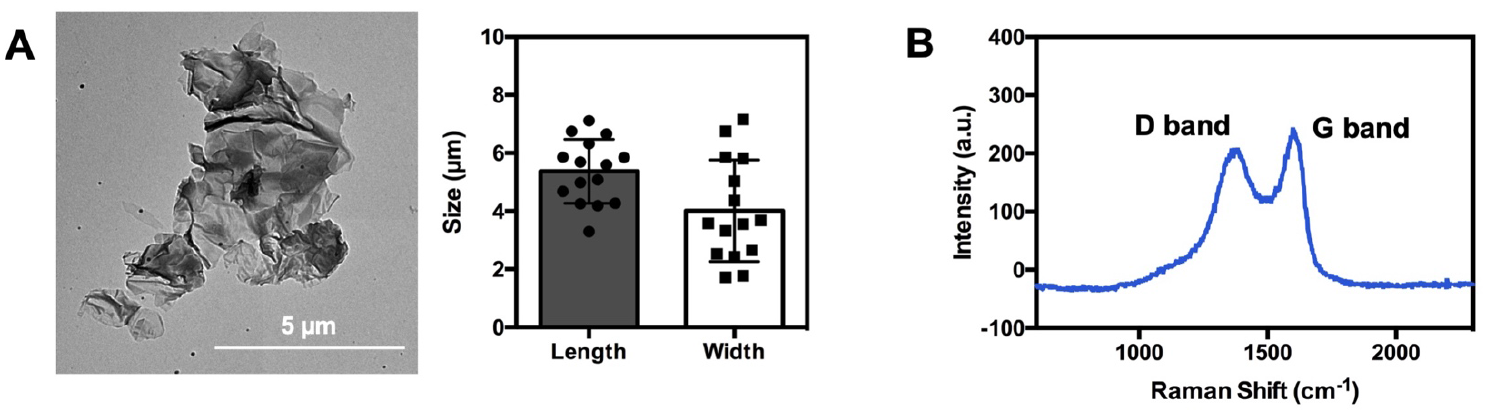
(A)Transmission electron microscopy image of an rGO flake and size analysis of flakes (n=15). (B) Raman spectra of the rGO flakes.

**Supporting Information Figure S2.**
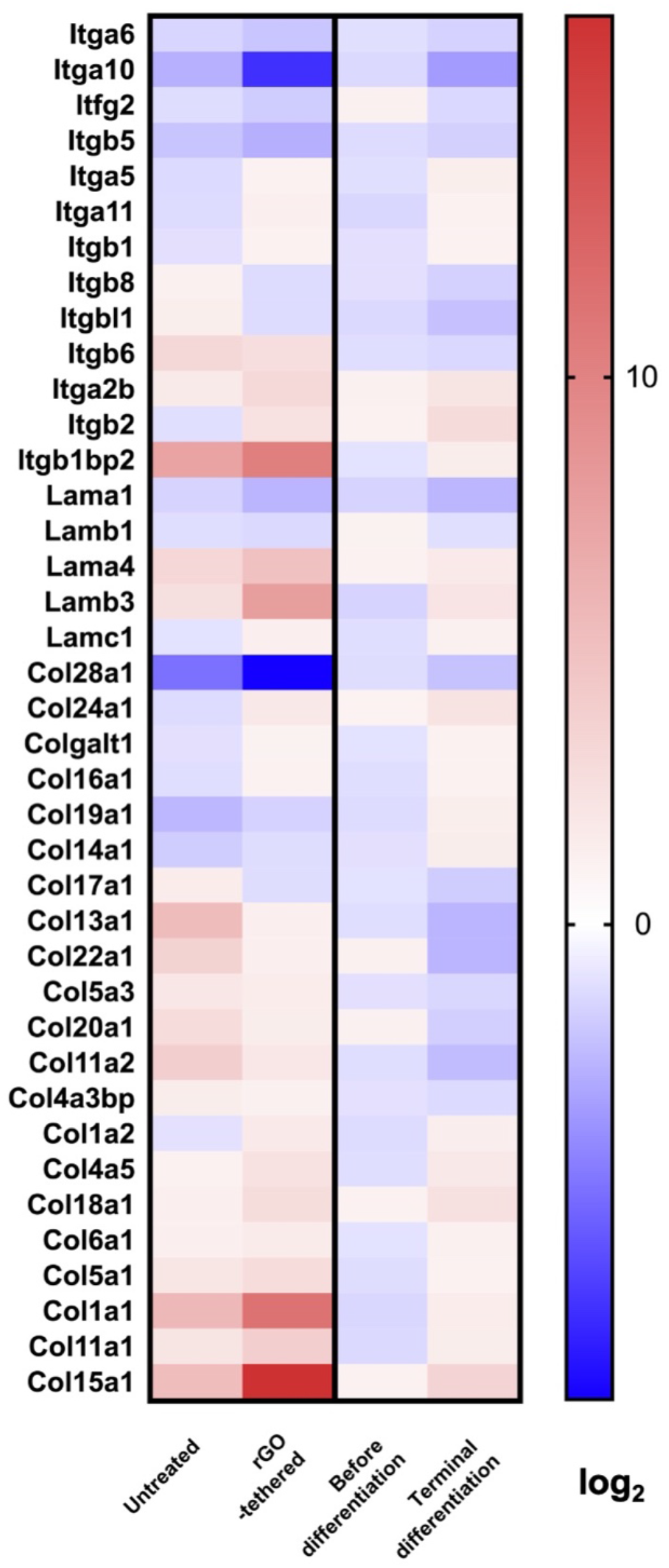
RNA sequencing profile of integrin, laminin, and collagen encoding genes. (FDR p-value<0.05).

